# Anticancer potential of soil-associated actinobacterium DHE 6-7 isolated from Enggano Island, Indonesia

**DOI:** 10.1101/2025.09.16.676699

**Authors:** Ika Nurzijah, Akhirta Atikana, Imas Amalia Wardani, Aditya Yuliandaru Pamungkas, Retno Wahyuningrum, Nunuk Aries Nurulita, Miranti Nurindah Sari, Shanti Ratnakomala, Fahrurozi Fahrurozi, Puspita Lisdiyanti

**Author notes:** Contributing authors. These authors contributed equally to this work (Ika Nurzijah, Akhirta Atikana).

## Abstract

The growing demand for pharmaceuticals has driven a shift towards developing medicinal products through bioprospecting by utilising sustainable sources. This involves exploring the potential of beneficial bacteria, such as Actinobacteria, known for producing diverse secondary metabolites with medicinal applications. In a previous study, 422 Actinobacteria were successfully isolated from Lombok, Bali, and Enggano Islands in Indonesia, and subjected to various bioactivity tests. Notably, Isolate DHE 6-7 from Enggano Island showed promising potential as a drug candidate. This study aimed to evaluate anticancer activity of DHE 6-7 isolates in T47D breast cancer cell lines and pinpoint the key metabolite contributing to its anticancer activity. The secondary metabolites from the DHE 6-7 culture were extracted using ethyl acetate by liquid-liquid extraction and its bioactive compound was analysed using thin layer chromatography. Subsequently, the extract underwent cytotoxic and antiproliferative assays in T47D cells. While Actinomycin-D was previously identified as a major bioactive compound in methanolic extract of DHE 6-7 isolate, this study confirmed that ethyl acetate extraction was able to retain the Actinomycin-D content from DHE 6-7 isolate. Interestingly however, the ethyl acetate extract of DHE 6-7 (EAE of DHE 6-7) showed superior cytotoxic activity in T47D cells compared to actinomycin-D alone. This may suggest the contribution of additional secondary metabolites in the EAE of DHE 6-7 to its anticancer activity. Notably, the combination of the EAE of DHE 6-7 isolate and 5-FU (5-fluorouracil) exhibited a synergistic effect, indicating the potential use of these compound(s) as a co-chemotherapeutic agent.

## 1 Introduction

Bioprospecting for drug discovery has recently shifted toward the exploration of sustainable biological resources [1–3]. This paradigm shift is driven by the increasing global demand for food and medicine, which has raised a critical dilemma: whether to prioritise natural resources for food security or as a source of bioactive compounds for therapeutic development [4]. At the same time, nature harbors a wealth of untapped potential, particularly among microorganisms. This realisation has sparked what has been termed as a “microbial revolution” in natural drug discovery [5–7].

Microorganisms have coexisted within Earth’s ecosystems for millions of years. Initially recognized as pathogens, the perception of microorganisms shifted dramatically following Alexander Fleming’s discovery of penicillin from *Penicillium notatum* in 1928. Since then, over 23,000 antimicrobial compounds have been identified, with more than two-thirds originating from actinobacteria [8, 9].

Actinobacteria are a diverse and ecologically significant phylum of Gram-positive bacteria, characterized by their high guanine and cytosine (G+C) content, typically exceeding 50%, and often reaching 70 mol% [10, 11]. They exhibit remarkable morphological differentiation, including the formation of substrate and aerial mycelia. Taxonomically, actinobacteria are broadly categorized into *Streptomyces* and non-*Streptomyces* groups. The *Streptomyces* genus is the most dominant and taxonomically diverse, encompassing hundreds of species known for producing extensive spore chains and powdery colonies. In contrast, non-*Streptomyces* actinomycetes, such as *Micromonospora, Actinoplanes, Nocardia*, and *Rhodococcus*, are less frequently isolated and often exhibit bacterial-like or compact morphologies [10, 12].

Beyond their role as prolific antimicrobial producers, actinobacteria synthesise a wide range of secondary metabolites with diverse applications, including anticancer, antifungal, antiparasitic agents, enzymes, and nanomaterials [8, 11, 13–17]. Remarkably, actinobacteria thrive in various ecological niches, including both terrestrial and aquatic environments [10, 12]. Motivated by Indonesia’s rich biodiversity, the Indonesian Culture Collection, National Research Center and Innovation Agency (BRIN) initiated a broad actinobacteria bioprospecting program targeting both soil and marine sources.

In 2015, as part of a nationwide actinomycete bioprospecting program, soil samples were collected from three diverse regions in Indonesia: Lombok, Bali, and Enggano Island [18]. This effort yielded a total of 422 actinomycetes isolated from the soil samples. Among them, an actinobacterium isolate DHE 6–7, obtained from Enggano Island, exhibited potential bioactivity against Gram-positive bacteria such as *Bacillus subtilis, Micrococcus luteus* and *Staphylococcus carnosus*.Subsequent molecular identification using 16S rRNA gene sequencing revealed a close phylogenetic relationship to *Streptomyces parvulus* NBRC 13193 (98.55% similarity). The species *Streptomyces parvulus* has been reported as producer of the polypeptide antibiotic actinomycin D [19, 20]. This compound is member of the actinomycin family, that is also notable for its potent dual activity as both antimicrobial and anticancer agents. Due to its high cytotoxicity, actinomycin D is primarily used in pediatric oncology rather than as a broad-spectrum antibiotic [21–23].

Given the rising global incidence of cancer, particularly breast cancer in women, the discovery of sustainable anticancer agents is critical for improving future therapeutic options. This study assessed the anticancer potential of the ethyl acetate extract (EAE) from isolate DHE 6-7, which showed strong cytotoxicity in T47D breast cancer cell lines. In addition to cytotoxic activity, antiproliferative assays and combination studies with 5-fluorouracil (5-FU) were conducted to evaluate synergistic effects. Although actinomycin D was confirmed as a constituent, its activity alone did not fully account for the bioactivity observed in the EEA of DHE 6–7, suggesting the involvement of additional bioactive metabolites.

## 2 Results

### 2.1 Rejuvenated colony of *Actinobacterium* DHE 6–7

Upon rejuvenation on ISP2 agar medium, the DHE 6–7 isolate demonstrated morphological characteristics typical of the genus *Streptomyces*. As shown in Figure 1, the colony formed a dense, powdery aerial mycelium with a whitish to pale-gray coloration, indicative of sporulation. The surface morphology was rough and irregular, which is characteristic of mature *Streptomyces* colonies.

**Fig. 1.**
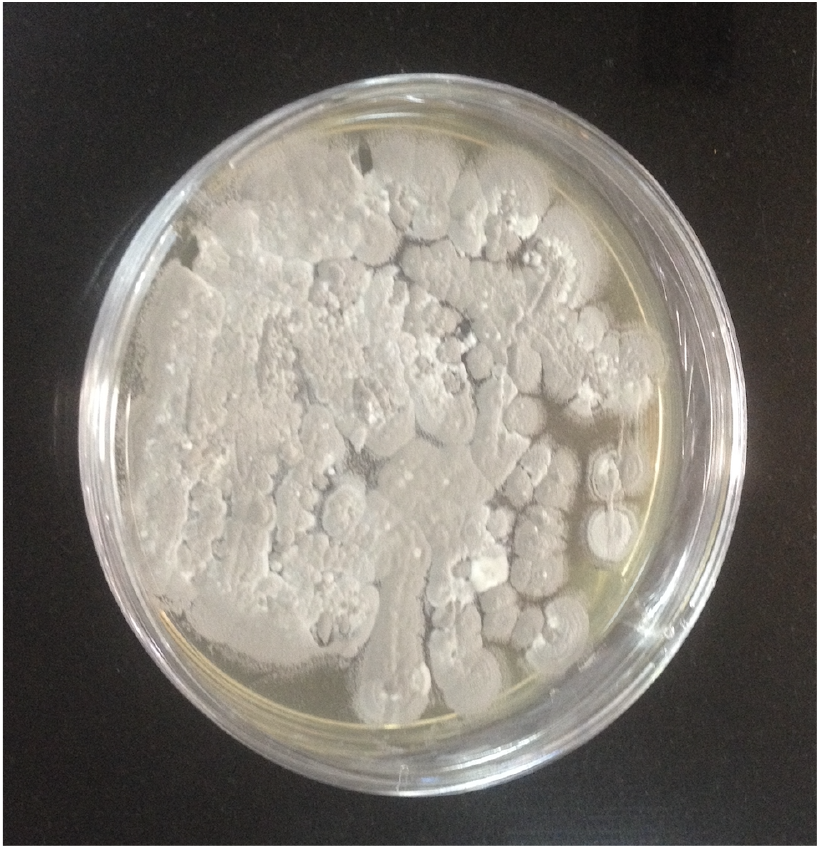
Rejuvenated colony of *Actinobacterium* DHE 6–7 on ISP2 agar medium. The culture displays typical *Streptomyces* morphology, with dense aerial mycelium and sporulation.

### 2.2 Characterisation of secondary metabolites by thin layer chromatography

Thin layer chromatography (TLC) was performed to characterise the secondary metabolite profile of the EAE of DHE 6–7. The TLC plate was developed using a mobile phase of hexane and ethyl acetate (0.5:9.5, v/v) and visualised under UV light at wavelengths of 254 nm and 366 nm. Actinomycin D was used as a reference solution. As shown in Figure 2, both the EAE of DHE 6–7 (lane A) and the actinomycin D standard (lane B) exhibited UV-active spots with comparable retention factor (Rf) values of 0.4 under 254 nm illumination. The co-migration of spots at 254 nm indicates the presence of compounds in EAE of DHE 6–7 with similar chromatographic profile to actinomycin D. Furthermore, weak fluorescent signals were observed under 366 nm UV light for the correponding compounds.

**Fig. 2.**
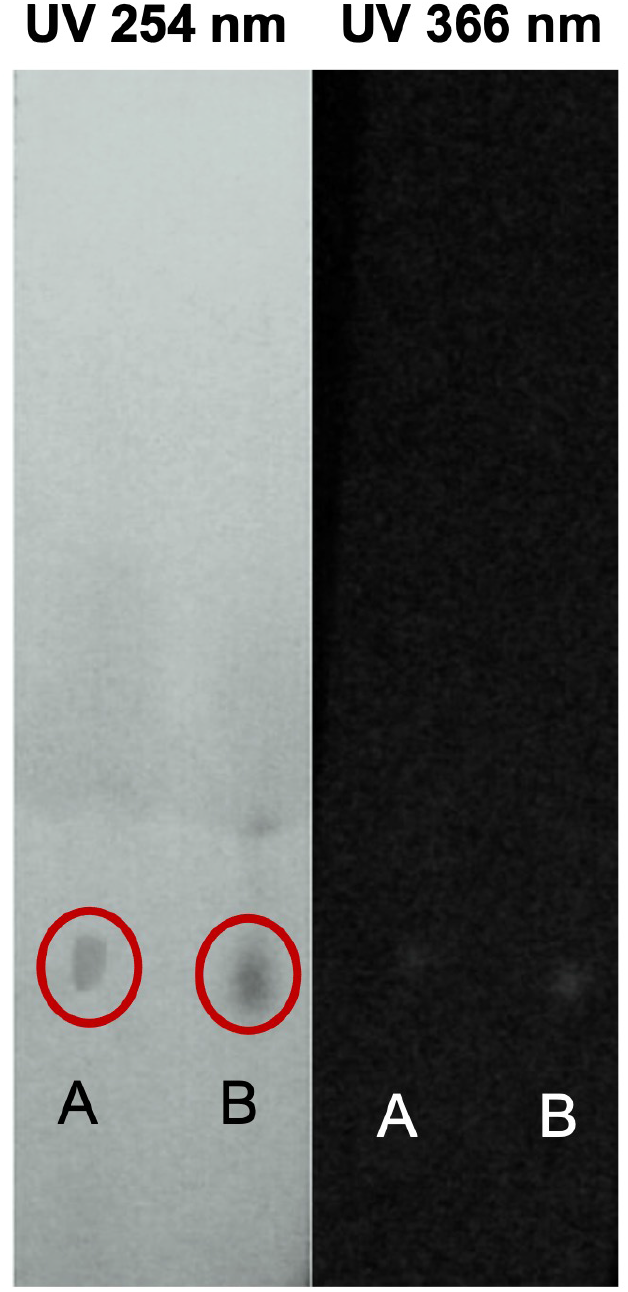
TLC plate visualised under UV 254 nm (left) and 366 nm (right). Lane A: EAE of DHE 6–7; Lane B: Actinomycin D reference solution. Red circles indicate the detected UV-active spots.

### 2.3 Cytotoxicity of ethyl acetate extract of DHE 6–7 in T47D cells

The cytotoxic effect of EAE of DHE 6–7 was evaluated in T47D breast cancer cells using the MTT assay after 24 hours of treatment. Cells were treated with serial concentrations of 0.01, 0.05, 0.1, 0.2, 0.4, 0.8, 1.0, and 2.0 *µ*g/mL of EAE of DHE 6–7. Morphological changes following treatment are shown in Figure 3, panel A. A progressive reduction in cell density and rounding of cells—typical of non-viable cells—was observed with increasing concentrations of the extract.

**Fig. 3.**
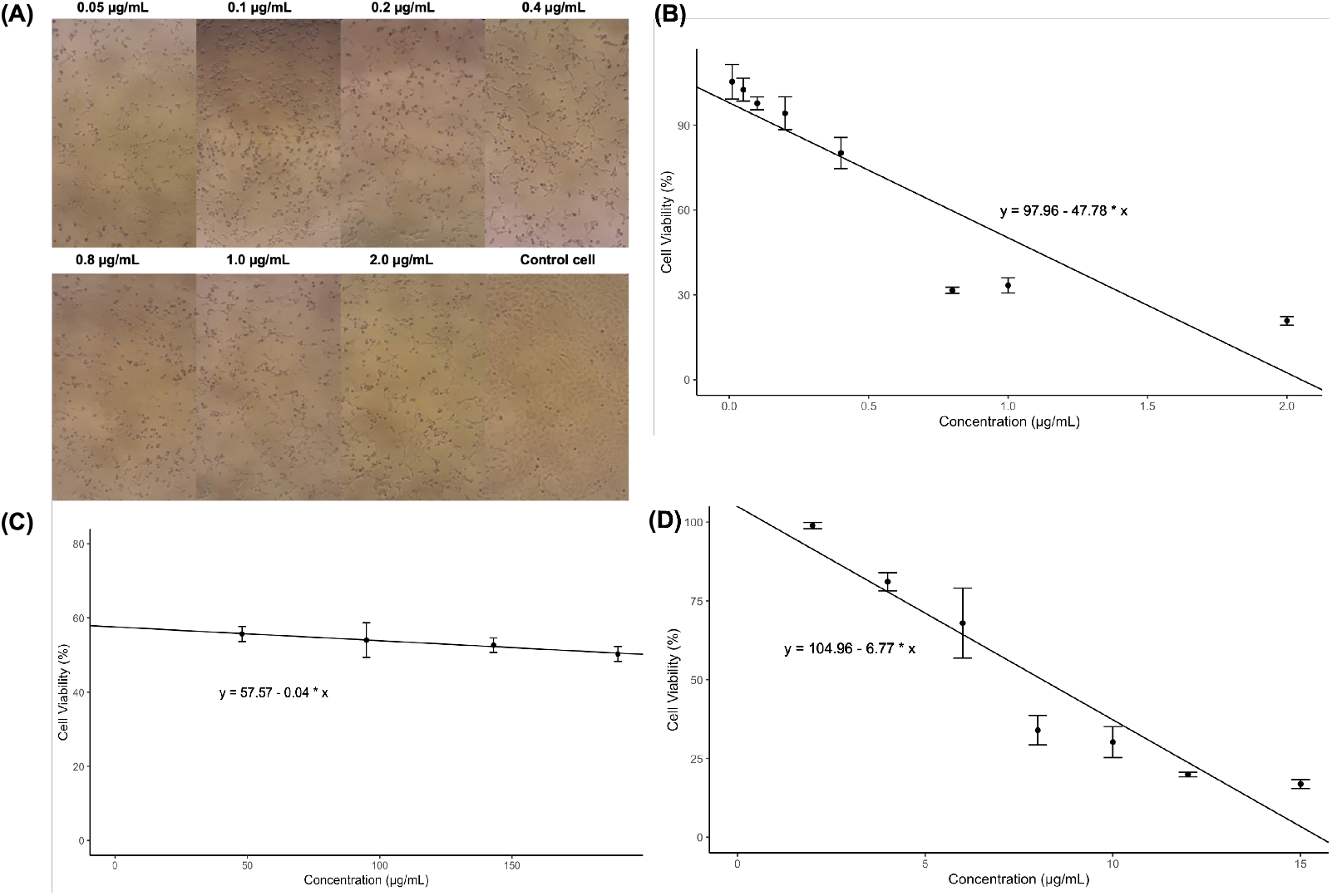
Cytotoxic effects of EAE of DHE 6–7, 5-FU, and actinomycin D in T47D cells after 24 hours of treatment. (A) Morphology of T47D cells after exposure to EAE of DHE 6–7 at increasing concentrations. (B) Dose–response curve of EAE of DHE 6–7 with IC_50_ = 1.00 *µ*g/mL. (C) Dose– response curve of 5-FU with IC_50_ = 203.93 *µ*g/mL. (D) Dose–response curve of actinomycin D with IC_50_ = 8.12 *µ*g/mL. Data represent mean *±* SD of triplicate experiments.

Dose–response curves were generated to determine the IC_50_ values of each compound (Figure 3, panels B–D) based on percentage cell viability. The IC_50_ value for EAE of DHE 6–7 was 1.00 *µ*g/mL. EAE of DHE 6–7 demonstrated greater cytotoxicity in T47D cells compared to the reference compound actinomycin D (IC_50_ = 8.12 *µ*g/mL) and the conventional chemotherapeutic agent 5-FU (IC_50_ = 203.93 *µ*g/mL). These results indicate that EAE of DHE 6–7 possesses higher anticancer potency than 5-FU and comparable activity to actinomycin D.

### 2.4 Cytotoxic combination of ethyl acetate extract of DHE 6–7 and 5-fluorouracyl

The combined cytotoxic effects of the EAE of DHE 6–7 and 5-FU in T47D cells were evaluated using the MTT assay. Combination index (CI) values were calculated for each treatment pair to assess the nature of the interaction. As shown in Figure 4, panel A, most combinations of EAE of DHE 6–7 with moderate to high concentrations of 5-FU (69–125 *µ*g/mL) yielded CI values greater than 1.0, indicating antagonistic effects. The strongest antagonism was observed at the highest combination doses, with CI values of 1.87 for 0.1 *µ*g/mL EAE of DHE 6–7 + 125 *µ*g/mL 5-FU and 1.86 for 0.1 *µ*g/mL EAE of DHE 6–7 + 94 *µ*g/mL 5-FU.

**Fig. 4.**
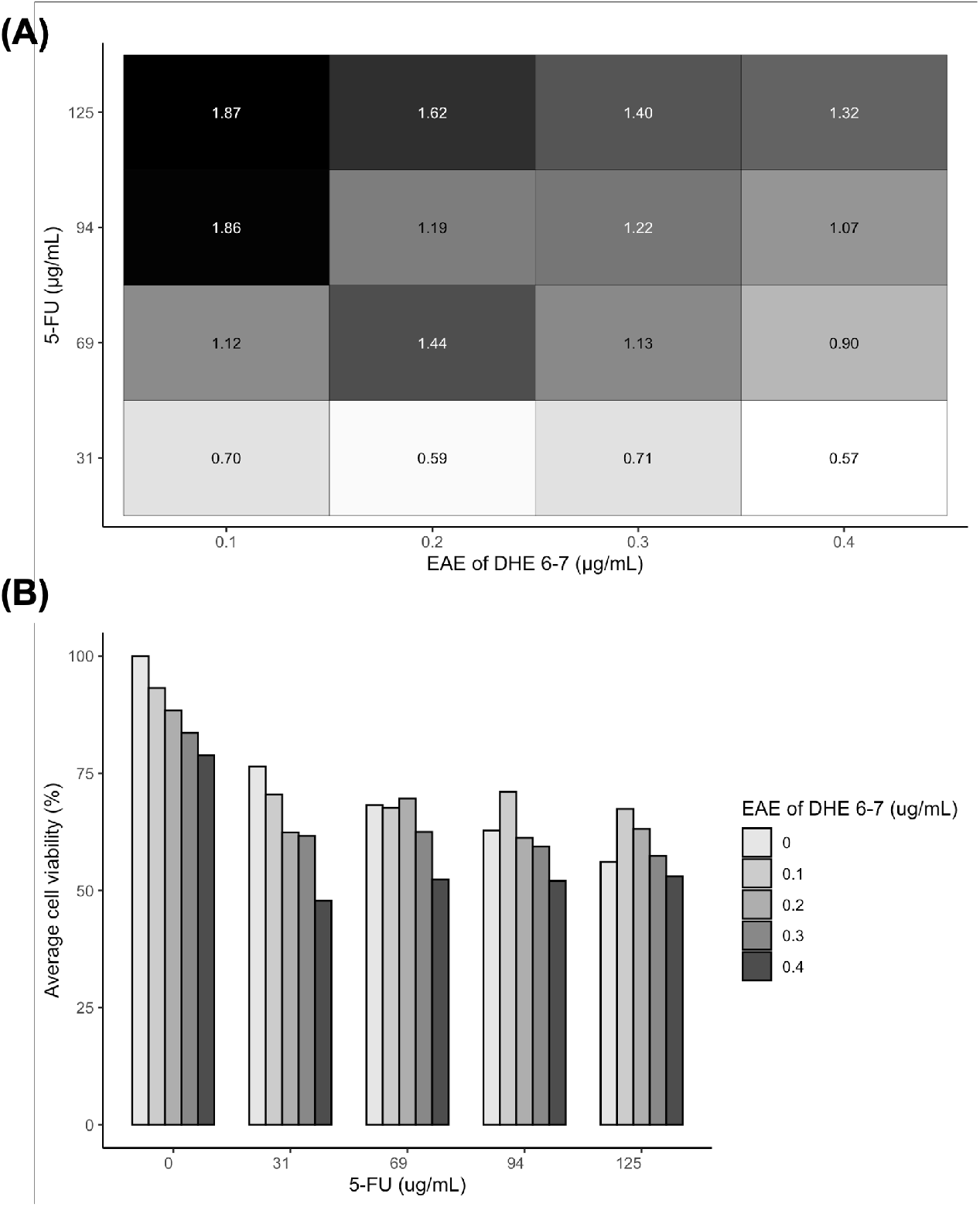
Cytotoxic combination of EAE of DHE 6–7 and 5-FU in T47D cells. (A) Heatmap of combination index (CI) values calculated from MTT assay data. CI values *<* 1 indicate synergism, CI = 1 additive, and CI *>* 1 antagonism. (B) Cell viability data (%) from 24-hour MTT assay at each combination dose. Values represent the mean of triplicate experiments.

In contrast, combinations involving all concentrations of EAE of DHE 6–7 (0.1–0.4 *µ*g/mL) with the lowest 5-FU dose (31 *µ*g/mL) produced CI values below 1.0, ranging from 0.57 to 0.71, indicative of synergistic effects. These findings suggest that low-dose 5-FU combinations enhance cytotoxic efficacy, while higher concentrations of 5-FU in combination result in diminished or antagonistic interactions.

This trend is further supported by the viability data (Figure 4, panel B). At 5-FU 31 *µ*g/mL, cell viability decreased progressively with increasing concentrations of EAE of DHE 6–7. The lowest viability (47.81%) was observed at the combination of 0.4 *µ*g/mL DHE and 31 *µ*g/mL 5-FU, confirming the synergistic interaction. In contrast, at higher 5-FU concentrations (94–125 *µ*g/mL), cell viability reductions were less pronounced and fluctuated with increasing concentrations of EAE of DHE 6–7, consistent with the observed antagonism.

### 2.5 Antiproliferative assay

The time-dependent antiproliferative effect of the EAE of DHE 6–7 was evaluated in T47D breast cancer cells over a 72-hour period using the MTT assay. As shown in Figure 5, panel A, treatment with EAE of DHE 6–7 at 0.2, 0.4, and 1.0 *µ*g/mL resulted in a fluctuating reduction in cell viability over time. While all doses caused an initial decline at 24 hours, a transient increase in viability was observed at 48 hours for some concentrations, followed by a further decrease at 72 hours. These results indicate that EAE of DHE 6–7 inhibits T47D cell proliferation, although the effect was not strictly dose- or time-dependent. For comparison, actinomycin D at 4, 8, and 20 *µ*g/mL induced a more pronounced and consistent reduction in viability, with values falling below 25% at 72 hours across all concentrations.

**Fig. 5.**
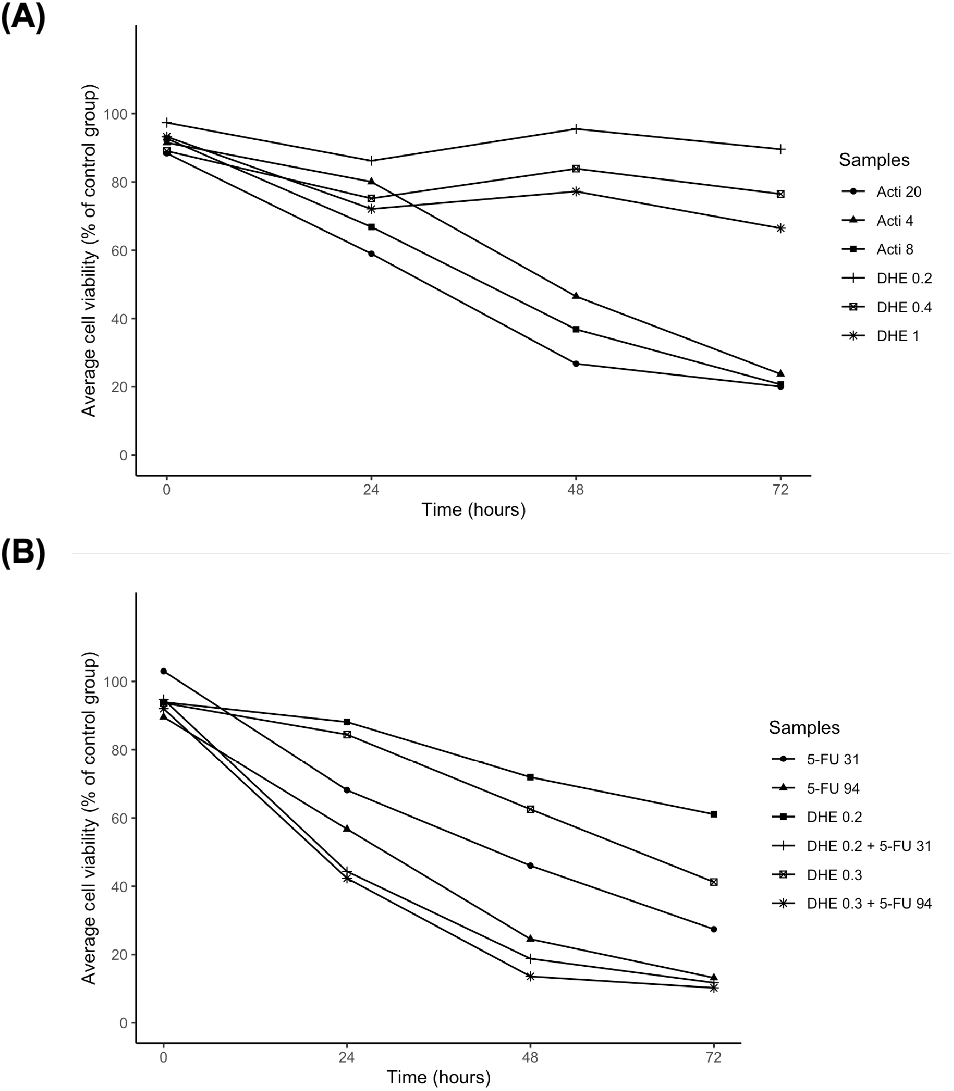
Antiproliferative activity of EAE of DHE 6-7 in T47D cells. (A) Time-course viability of T47D cells treated with actinomycin D (4, 8, and 20 *µ*g/mL) or EAE of DHE 6-7 (0.2, 0.4, and 1.0 *µ*g/mL). (B) Time-course viability of T47D cells treated with 5-FU (31 and 94 *µ*g/mL) alone or in combination with EAE of DHE 6-7 (0.2 and 0.3 *µ*g/mL). Cell viability was assessed at 0, 24, 48, and 72 hours using the MTT assay. Values represent the mean of triplicate experiments.

To assess the antiproliferative potential of the combination of EAE of DHE 6–7 with 5-FU, selected concentrations of both agents were co-administered and monitored across the same time points (Figure 5, panel B). Combinations of 0.2 *µ*g/mL EAE of DHE 6–7 (1/5 IC_50_) with 31 *µ*g/mL 5-FU and 0.3 *µ*g/mL EAE of DHE 6–7 (1/3 IC_50_) with 94 *µ*g/mL 5-FU resulted in greater suppression of viability than either agent alone. The most pronounced effect was observed with the 0.3 *µ*g/mL DHE + 94 *µ*g/mL 5-FU combination, which reduced cell viability to 10.24% at 72 hours, suggesting an enhanced antiproliferative effect.

## 3 Materials and Methods

### 3.1 Rejuvenation of Actinobacterium DHE 6-7

The Actinobacterium DHE 6-7 was isolated from soil samples of Enggano Island (5°2257.08” S, 102°1328.28” E), Indonesia. The soil samples was collected at a depth of approximately 10 cm, in December 2015. The cultivation of actinomycetes isolates from soil samples of Enggano was performed using a serial dilution technique on Humic Acid–Vitamin (HV) agar medium, following a dry heat pretreatment, as described in published paper [18]. The selected isolate DHE 6-7 was maintained in the Streptomyces International Project 2 (ISP2) agar medium for morphological characterization and molecular identification as described in published paper [18].

### 3.2 Extraction of secondary metabolites

The isolate DHE 6–7 was subcultured on R5 agar medium [24] and incubated at 28 °C for 7 days. A single colony was then inoculated into 10 mL of tryptic soy broth (TSB) and incubated at 28 °C with shaking at 180 rpm for 2 days. Subsequently, 2 mL of this pre-culture was transferred into 30 mL of R5 liquid production medium in a 250 mL baffled Erlenmeyer flask. The R5 production medium consisted of sucrose (103 g/L), glucose (10 g/L), KSO (0.25 g/L), MgCl (10.12 g/L), casamino acids (0.1 g/L), yeast extract (5 g/L), and 2 mL/L of trace element solution (TES), with the pH adjusted to 7.2. Cultivation was carried out at 28 °C with agitation at 180 rpm for 4 days. After incubation, the culture broth was centrifuged at 3000 rpm for 5 minutes to separate the biomass. The resulting supernatant was extracted with ethyl acetate (1:1, v/v; Sigma-Aldrich, USA). The organic phase was evaporated using a rotary evaporator (IKA RV–8) at 40 °C, and the residue was reconstituted in 500 µL of methanol for further analysis.

### 3.3 Thin layer chromatography (TLC)

Thin layer chromatography (TLC) was performed to characterise the secondary metabolites present in EAE of DHE 6–7. Silica gel GF254 plates (analytical grade) were used as the stationary phase and pre-activated at 110°C for 30 minutes. The mobile phase consisted of hexane and ethyl acetate (0.5:9.5, v/v). Aliquots (1 *µ*L) of both the EAE of DHE 6–7 and a reference solution of actinomycin D (Thermo Fisher Scientific, UK) diluted in DMSO were applied to the plate and developed in a saturated TLC chamber. After elution, the plates were examined under ultraviolet (UV) light at 254 nm and 366 nm to detect UV-active compounds. Retention factor (Rf) values were calculated for each visible spot and compared to the standard to evaluate the possible presence of actinomycin D in EAE of DHE 6–7. The Rf value was determined using the following equation:

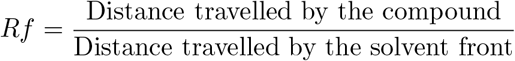

### 3.4 Cytotoxicity assay

The cytotoxic activity of EAE of DHE 6-7, actinomycin D, and 5-fluorouracil (5-FU; Merck, Germany) was assessed in T47D breast cancer cells using the MTT assay. Cells were seeded into 96-well plates at a density of 5 × 10^3^ cells/well and incubated for 24 hours at 37°C in a humidified atmosphere with 5% CO_2_ to allow attachment and recovery. Dulbecco’s Modified Eagle Medium (DMEM; Gibco, Thermo Fisher Scientific, UK), supplemented with 10% fetal bovine serum (FBS) and 1% penicillin-streptomycin, was used as the culture medium.

Test compounds were initially dissolved in dimethyl sulfoxide (DMSO; Merck, Germany) to prepare stock solutions. The EAE of DHE 6-7 was prepared at 1000 µg/mL and serially diluted in DMEM to final concentrations of 0.01, 0.05, 0.1, 0.2, 0.4, 0.8, 1.0, and 2.0 µg/mL. Actinomycin D was tested at concentrations of 2, 4, 6, 8, 10, 12, and 15 µg/mL, while 5-FU was evaluated at 48, 95, 143, and 190 µg/mL. The final concentration of DMSO in the culture medium did not exceed 1.5%.

After incubation, the media were discarded and the cells were washed with phosphate-buffered saline (PBS). Then, 100 µL of MTT solution (0.5 mg/mL in DMEM) was added to each well and incubated for 4 hours. Following incubation, 100 µL of 10% SDS in 0.1 N HCl was added to solubilise the formazan crystals, and the plates were incubated overnight at room temperature in the dark. Absorbance was measured at 595 nm using a microplate reader. Cell viability was calculated relative to the untreated control, and the IC_50_ values were determined by nonlinear regression analysis using using R Studio for macOS (version 2024.12.1) [25].

### 3.5 Antiproliferative activity

The time-dependent cytotoxic effects of the EAE of DHE 6-7, actinomycin D, and 5-FU were evaluated by measuring cell viability over a 72-hour period. T47D cells were seeded into 96-well plates at a density of 2.5 × 10^3^ cells/well and incubated for 24 hours to allow for attachment. Cells were then treated with EAE of DHE 6–7 (0.2, 0.3, and 1 µg/mL), actinomycin D (4, 8, and 20 µg/mL), and 5-FU (31 and 94 µg/mL), either individually or in combination. Treatments were performed in triplicate.

Cell viability was assessed at 0, 24, 48, and 72 hours using the MTT assay, as described previously. Absorbance readings at 595 nm were recorded at each time point and normalised to the control group to calculate the percentage of viable cells. This time-course analysis provided a quantitative assessment of cytotoxic progression for each treatment group, with decreasing viability indicating time-dependent cytotoxic or antiproliferative effects.

### 3.6 Cytotoxicity of combination treatment

The cytotoxic potential of the combination of ethyl acetate extract of DHE 6-7 (EAE of DHE 6-7) and 5-fluorouracil (5-FU) was evaluated in T47D breast cancer cells using the MTT assay at a single time point. Cells were seeded into 96-well plates at a density of 5 × 10^3^ cells/well and incubated for 24 °C in a 5% CO_2_ incubator for 24 hours to allow attachment. Cells were then treated with EAE of DHE 6–7 alone, 5-FU alone, or combinations of both agents at the following concentration pairs: (1) 0.2 *µ*g/mL DHE + 31 *µ*g/mL 5-FU, (2) 0.3 *µ*g/mL DHE + 94 *µ*g/mL 5-FU.

All treatments were performed in triplicate. After 24 hours of exposure, cell viability was assessed using the MTT assay, as previously described. Absorbance was measured at 595 nm, and the percentage of viable cells was calculated relative to untreated controls.

The interaction between EAE of DHE 6-7 and 5-FU was evaluated using the Combination Index (CI), following the Chou–Talalay method [26]. CI values were calculated in R Studio using experimentally derived viability data and the following formula:

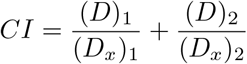

where (*D*)_1_ and (*D*)_2_ represent the concentrations of each drug in the combination required to produce an *x*% effect, and (*D*_*x*_)_1_ and (*D*_*x*_)_2_ represent the concentrations of the individual agents needed to achieve the same effect alone. A CI value *<* 1 indicates synergism, CI = 1 denotes an additive effect, and CI *>* 1 indicates antagonism.

## 4 Discussion

The current study applies bioprospecting of anticancer agents using sustainable resources such as actinobacteria. This was achieved by investigating the anticancer potential of an ethyl acetate extract derived from the actinobacterium isolate DHE 6–7 (EAE of DHE 6–7), focusing on its cytotoxic and antiproliferative effects in T47D breast cancer cells. The rejuvenated DHE 6–7 grown on ISP2 agar medium exhibited *Streptomyces*-like characteristics, with colonies displaying powdery aerial mycelium and a whitish to pale-gray coloration, suggestive of sporulation [10–12]. These findings support a previous study that identified DHE 6–7 as *Streptomyces parvulus* NBRC 13193 (98.55% similarity) based on 16S rRNA sequencing [18].

The TLC analysis demonstrated that the extract contained UV-active compounds with similar Rf values to the reference standard actinomycin D, suggesting the presence of structurally related metabolites. The potential of isolate DHE 6-7 as producer of actinomycin D has also been confirmed by the HPLC/MS analysis of its methanolic extracts [18]. Although the exact constituents in EAE of DHE 6-7 need confirmation, cytotoxic activity in EAE of DHE 6-7 was probably due to the actinomycin D in the extracts. These findings support prior studies on the potential of actinobacteria as prolific producers of secondary metabolites with anticancer activity [15, 27].

In the MTT cytotoxicity assay, the EAE of DHE 6–7 showed a strong inhibitory effect on the viability of T47D breast cancer cells, with an IC_50_ value of 1.00 *µ*g/mL. This potency was markedly higher than that of the standard chemotherapeutic agent 5-FU, which had an IC_50_ of 203.93 *µ*g/mL, and even surpassed the activity of actinomycin D (IC_50_ = 8.12 *µ*g/mL). Morphological changes and a visible reduction in cell density observed following treatment with serial concentrations of EAE DHE 6–7 further confirmed its cytotoxic activity.

These findings are consistent with previous studies on the anticancer properties of both 5-FU and actinomycin D. A study reported that 5-FU had an IC_50_ of 213 ± 2.2 *µ*g/mL against T47D breast cancer cells, which aligns with our current findings [28]. A purified actinomycin D from *Streptomyces parvulus* isolated from a mangrove ecosystem in Kerala, India, exhibited potent cytotoxic activity against A459 lung cancer cells, with an IC_50_ value of 0.52 *µ*g/mL [20]. In a follow-up study, actinomycin D also demonstrated IC_50_ values of 0.90 *µ*g/mL against U251 (glioblastoma), 1.09 *µ*g/mL against HCT-116 (colon cancer), and 0.56 *µ*g/mL against MCF-7 (breast cancer) after 48 hours of incubation. Additional studies demonstrated the dose- and time-dependent cytotoxicity of actinomycin D, with its most potent effect observed against U251 cells, where the IC_50_ decreased from 1.07 *µ*g/mL at 24 hours to 0.56 *µ*g/mL at 48 hours and 0.028 *µ*g/mL at 72 hours. Morphological changes in U251 cells treated with actinomycin D were also confirmed by acridine orange and ethidium bromide staining, indicating apoptotic activity [29].

In contrast, our study showed that despite the presence of actinomycin D in the EAE of DHE 6–7, it exhibited relatively lower cytotoxicity in T47D cells (IC_50_ = 8.12 *µ*g/mL) compared to the values reported for other cancer cell lines. This variation may be due to differences in cell line sensitivity. Notably, the extract as a whole displayed much stronger cytotoxicity (IC_50_ = 1.00 *µ*g/mL), suggesting that additional secondary metabolites in EAE DHE 6–7—beyond actinomycin D—may contribute significantly to the anticancer effect. These results support the hypothesis that unidentified bioactive compounds within the extract could be acting synergistically or independently to enhance its cytotoxic activity in T47D cells.

The combination treatment study revealed important insights into dose-dependent interaction behavior. While most combinations of EAE of DHE 6–7 with moderate to high concentrations of 5-FU resulted in antagonistic effects (CI *>* 1), all combinations involving low-dose 5-FU (31 *µ*g/mL) and increasing concentrations of the extract showed CI values below 1.0, indicating synergism. These findings suggest that at lower doses, EAE of DHE 6–7 may enhance the anticancer effect of 5-FU, allowing the drug to work more effectively than it would alone. This supports the potential role of EAE of DHE 6–7 as a co-chemotherapeutic agent, consistent with emerging strategies that combine natural products with conventional drugs to improve therapeutic outcomes and reduce toxicity [30–33].

The antiproliferative time-course assay revealed that EAE of DHE 6–7 showed no clear dose- or time-dependent effect. A transient increase in cell viability was observed at 48 hours, followed by a modest decline at 72 hours. Despite its strong cytotoxic activity, EAE of DHE 6–7 alone failed to demonstrate a consistent antiproliferative effect in T47D cells. However, when combined with 5-FU, particularly at 0.3 *µ*g/mL of the extract and 94 *µ*g/mL of 5-FU, a significant enhancement in growth inhibition was observed, with cell viability reduced to just over 10%. These results indicate that EAE of DHE 6–7 may enhance the antiproliferative activity of 5-FU in T47D cells. Further investigation is needed to clarify the molecular mechanisms involved and to identify other pathways that may contribute to the anticancer activity of EAE of DHE 6–7 in T47D cells.

Collectively, these findings highlight the potential of actinobacterium DHE 6–7 as a promising source of anticancer compounds. However, further isolation, purification, and identification of the individual bioactive constituents are important, especially to identify and to elucidate their mechanisms of action. Future work should also assess the selectivity of EAE of DHE 6–7 toward cancer cells versus normal cells and validate its efficacy using in vivo models. In addition, genomic approaches such as whole genome sequencing of the isolate DHE 6-7 also essential to assess the biosynthetic gene cluster, especially to elucidate pathways for production of target compounds. For example, to leverage the production of actinomycin D from actinobacterium DHE 6-7.

## 5 Conclusion

This study highlights the potent cytotoxic activity of the ethyl acetate extract from actinobacterium DHE 6–7 against T47D breast cancer cells. The extract showed stronger effects than 5-FU and actinomycin D, and synergistically enhanced 5-FU activity at low doses. Although its antiproliferative pattern was variable, the overall findings suggest its potential as a co-chemotherapeutic agent.

Future studies should focus on isolating and characterizing the secondary metabolites of DHE 6–7, evaluating their selectivity and *in vivo* efficacy, and employing genomic approaches such as whole-genome sequencing to identify biosynthetic gene clusters involved in secondary metabolite production, including actinomycin D.

